# Comparison of Cosine, Modified Cosine, and Neutral Loss Based Spectrum Alignment For Discovery of Structurally Related Molecules

**DOI:** 10.1101/2022.06.01.494370

**Authors:** Wout Bittremieux, Robin Schmid, Florian Huber, Justin JJ van der Hooft, Mingxun Wang, Pieter C Dorrestein

## Abstract

Spectrum alignment of tandem mass spectrometry (MS/MS) data using the modified cosine similarity and subsequent visualization as molecular networks have been demonstrated to be a useful strategy to discover analogs of molecules from untargeted MS/MS-based metabolomics experiments. Recently, a neutral loss matching approach has been introduced as an alternative to MS/MS-based molecular networking, with an implied performance advantage in finding analogs that cannot be discovered using existing MS/MS spectrum alignment strategies. To comprehensively evaluate the scoring properties of neutral loss matching, the cosine similarity, and the modified cosine similarity, similarity measures of 955,228 peptide MS/MS spectrum pairs and 10 million small molecule MS/MS spectrum pairs were compared. This comparative analysis revealed that the modified cosine similarity outperformed neutral loss matching and the cosine similarity in all cases. The data further indicated that the performance of MS/MS spectrum alignment depends on the location and type of the modification, as well as the chemical compound class of fragmented molecules.

## Introduction

With the growing availability of free, open, and proprietary tandem mass spectrometry (MS/MS) libraries, MS/MS-based spectrum alignment is routinely used for metabolite annotation and organization of mass spectral datasets.^1–4^ For MS/MS spectrum comparison, the cosine similarity is the most widely used approach to match near-identical spectra with each other.^5,6^ Adaptations hereof, such as the modified cosine similarity,^1,7^ can be used to discover non-identical but related spectra using analog searching and molecular networking of small molecules.^8,9^ Similar approaches have also been used in proteomics to perform open modification searching for the unbiased detection of peptides that contain post-translational modifications.^10–13^ Additionally, spurred by the emergence of increasingly powerful computational and machine learning techniques, several novel similarity scores have been proposed.^14–18^

Recently, neutral loss-based spectrum alignment was introduced in the METLIN analysis ecosystem^19^ as a strategy to match MS/MS spectra of analog molecules.^20^ During neutral loss matching, MS/MS spectra are mirrored at their precursor mass by calculating the distances from each fragment ion peak to the precursor mass, describing the neutral losses. Effectively, the neutral loss approach recalibrates spectra with their precursor masses as the origin, and a mirror database of neutral loss spectra was created in METLIN.^19^ The neutral loss spectra can then be used to find related spectra of modified analog molecules. Based on representative examples, Aisporna et al.^20^ have postulated that neutral loss matching can be particularly effective at finding MS/MS spectrum pairs that other approaches, such as spectrum alignment using the cosine and modified cosine similarities, could not find. This benefit was explained by the increase in spectrum usage (i.e. more explained intensity captured by matching fragment ions) in the assignment of related matches. Nevertheless, although a few select MS/MS spectra were chosen to compare different spectrum similarity measures, the difference in spectrum matching performance was not systematically quantified.^20^

We hypothesized that the modified cosine similarity, which forms the foundation of molecular networking in the Global Natural Products Social Molecular Networking (GNPS) system,^7^ should be able to discover all MS/MS spectrum pairs that can be found by neutral loss matching, as it considers both directly matching peaks and matching peaks that are shifted by the pairwise precursor mass difference. We, therefore, set out to systematically compare the performance of neutral loss matching, cosine similarity, and modified cosine similarity on a large collection of spectrum pairs of modified peptides and small molecules. As the data of the original study^20^ was not available due to the access restrictions of the METLIN database,^19^ we instead used a large collection of 955,228 and 10 million MS/MS spectrum pairs for peptides and small molecules, respectively, which were derived from the MassIVE-KB^21^ and GNPS resources.^7^

## Results

We evaluated neutral loss matching, cosine similarity, and modified cosine similarity in matching MS/MS spectra of non-identical yet structurally related molecules (**Figure 1**). The cosine similarity is a common strategy to compare MS/MS spectra to each other by matching fragment ions in both spectra with identical *m/z* values (while accounting for a relevant fragment mass tolerance based on the data acquisition settings). The modified cosine similarity is an extension of the cosine similarity which not only matches fragment ions with identical *m/z* values, but also considers fragment ions across both spectra with *m/z* values that are shifted by the precursor mass difference of the spectrum pair under consideration. Finally, neutral loss matching first transforms MS/MS spectra into neutral loss spectra by mirroring the fragment ion *m/z* values at the precursor mass, after which the neutral loss peaks with identical Δ*m/z* values are matched to each other.

**Figure 1.**
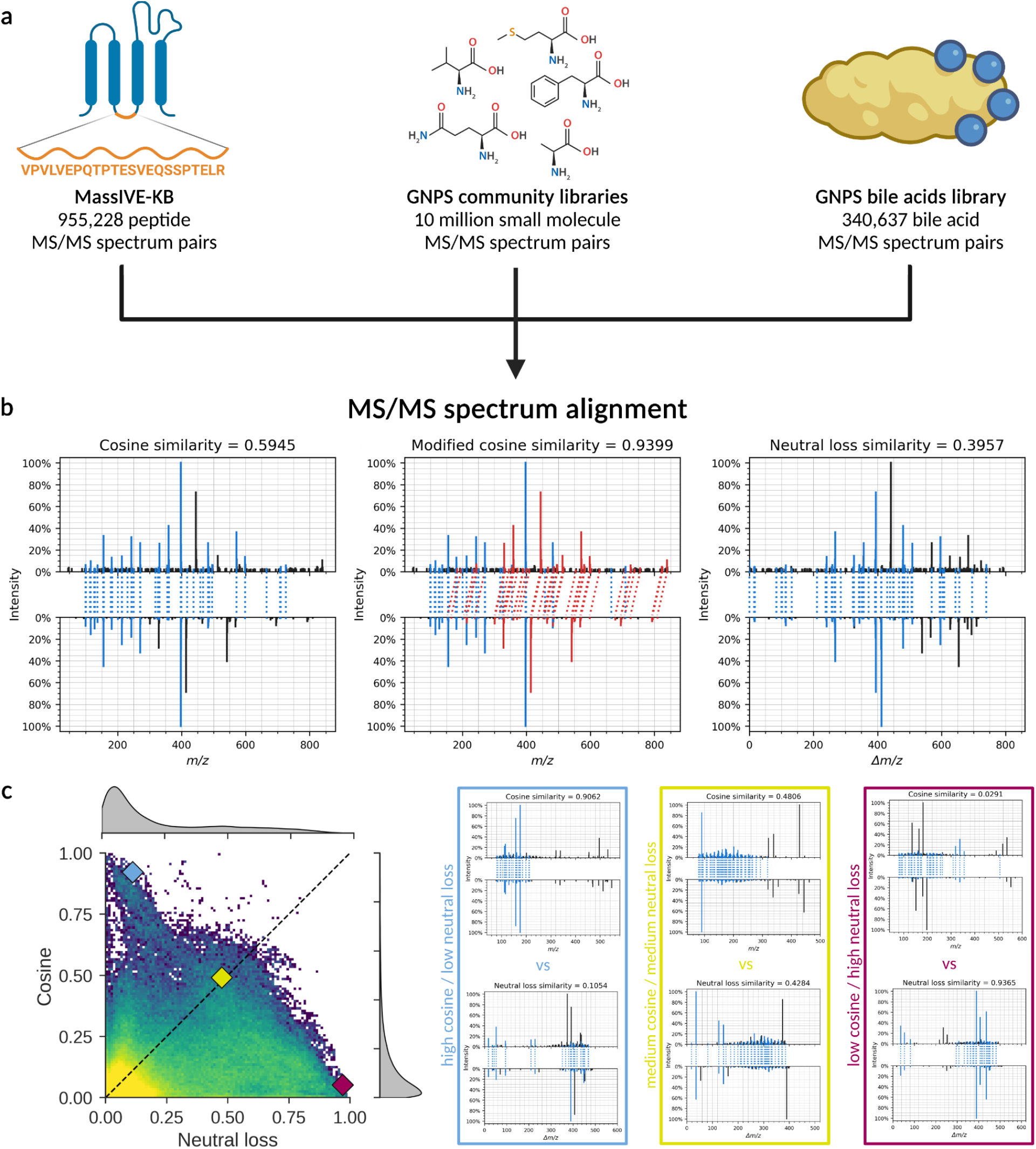
Evaluation of MS/MS spectrum similarity measures. (**a**) Spectrum similarity measures are evaluated using large MS/MS datasets drawn from the MassIVE-KB peptide spectral library, the small molecule GNPS community spectral libraries, and the GNPS bile acids library. (**b**) Illustration of the MS/MS spectrum alignment methods applied to the natural product apratoxin A (top; precursor *m/z* = 840.497) and a newly discovered apratoxin analog (bottom; precursor *m/z* = 810.487).^22^ We evaluate the cosine similarity (left), modified cosine similarity (middle), and neutral loss matching (right, visualized as neutral loss spectra) to compare MS/MS spectra to each other. Directly matching fragments with near-identical *m/z* or Δ*m/z* values in the original MS/MS spectra and the neutral loss spectra, respectively, are indicated in blue. Additional matching fragments that are offset by the pairwise precursor *m/z* difference, considered by the modified cosine similarity, are indicated in red. Unmatched fragments are indicated in black. (**c**) Illustration of a heat map to compare two alternative spectrum similarity measures, specifically the cosine similarity and neutral loss matching. The heat map shows the bivariate distribution of the corresponding spectrum similarity values for all MS/MS spectrum pairs, with each point corresponding to multiple observations as indicated by the heat map coloring. The density plots (gray) show the marginal distributions of the individual spectrum similarities. Representative examples with different spectrum similarity values are indicated by the colored diamonds, with the corresponding spectrum alignments shown in the colored boxes.

Three datasets with heterogeneous mass spectral properties were used to compare the different spectrum similarity measures. Because large-scale ground truth benchmark datasets of related small molecule spectrum pairs with known structural modifications currently do not exist yet, the first dataset we used consisted of 955,228 known peptide MS/MS spectrum pairs derived from the MassIVE-KB resource,^21^ where peptide pairs differ by a single modification (post-translational modification or amino acid substitution). Second, mass spectrum pairs were derived from 495,600 small molecule reference MS/MS spectra contained in the GNPS community mass spectral libraries.^7^ Because over 1.5 billion pairwise comparisons (precursor mass difference between 1–200 Da) can be performed using these spectra, a random subset of 10 million spectrum pairs was used for computational efficiency. Third, 340,637 bile acid MS/MS spectrum pairs from the GNPS community spectral libraries were used to evaluate the influence of specific modification properties, such as addition or loss of oxygen, conjugation of the bile acids with amino acids (as amino acid amidates), and substitution of conjugated amino acids attached to the bile acid core.

Evaluation of the different spectrum similarity measures on the peptide data indicates that the modified cosine similarity strictly outperforms both the cosine similarity and neutral loss matching (**Figure 2a**), while neutral loss matching resulted in a higher score than the cosine similarity in only 8.8% of the peptide pairs. Spectrum usage (in terms of the number of explained fragment ions and total explained intensity) can be used to assess the performance of spectrum similarity measures (**Figure 2b**).^20^ In general, neutral loss matching resulted in very low spectrum usage for peptide mass spectra, with a median explained intensity of 17.1%. In contrast, the median explained intensity for the cosine similarity and modified cosine similarity were 56.4% and 70.7%, respectively. Taking into account the modification position relative to the linear peptide sequences (**Figure 2c**), neutral loss matching performed better with modifications located closer to the C-terminal end. In this case, a maximal number of y-ions, which are the dominant signals in peptide mass spectra, include the modification and can be matched using the neutral loss strategy. In contrast, the cosine similarity performed best when the modification was located close, but not entirely at the peptide N-termini. We hypothesize that in this case the cosine similarity can match a large number of y-ions as well as the initial b-ions, whereas when the modification occurs at the N-terminus, no b-ions contribute to the cosine similarity while higher index y-ions often do not occur and thus do not provide further benefit. Finally, the modified cosine similarity achieves high scores across a range of modification positions (except for N-terminal modifications), which indicates its beneficial performance in capturing both directly matching fragment ions and fragment ions that are shifted due to the mass difference induced by the modification.

**Figure 2.**
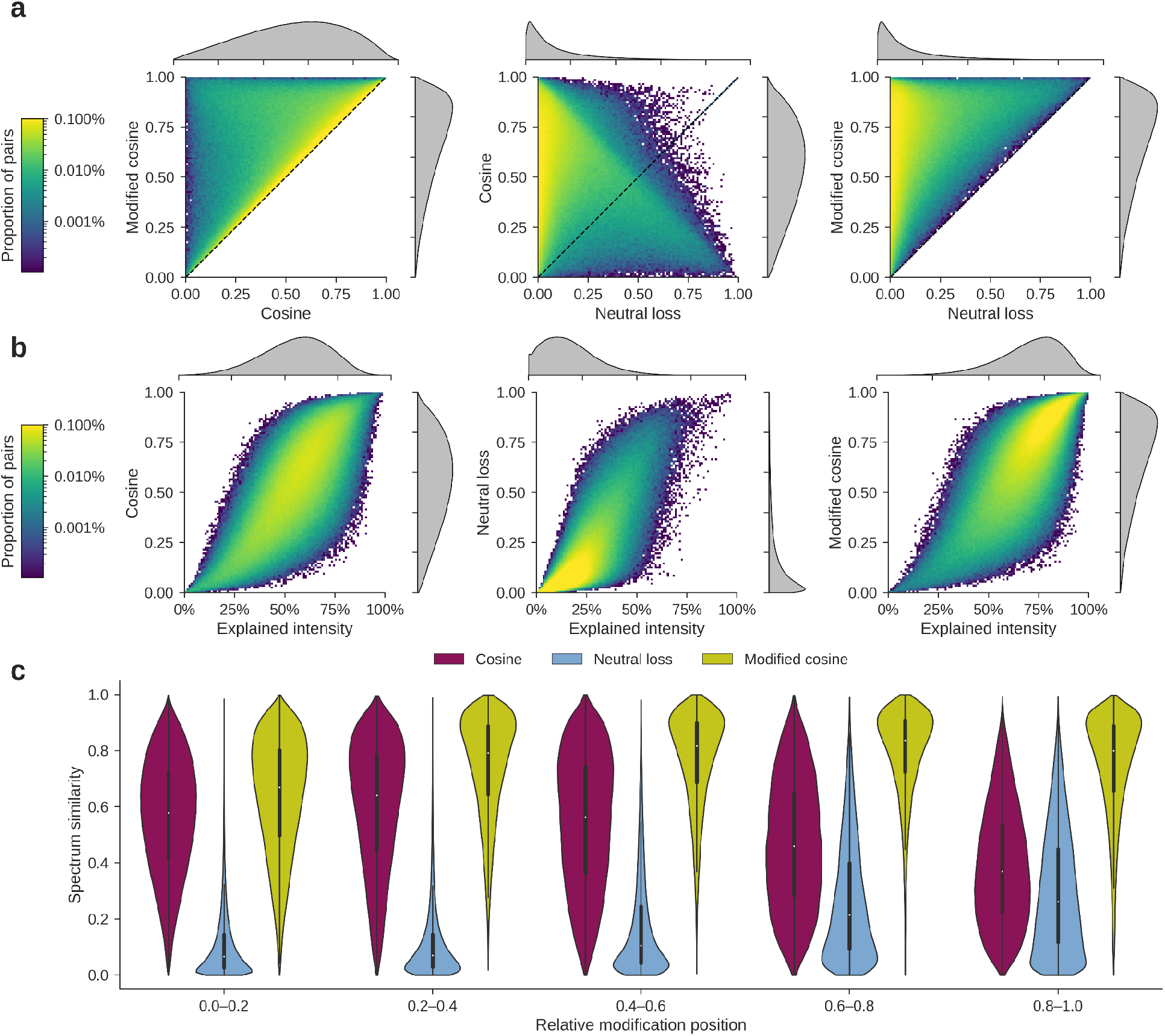
Comparison between the cosine similarity, neutral loss matching, and the modified cosine similarity when matching modified peptide MS/MS spectra. (**a**) Comparison between cosine similarity and modified cosine similarity (left), neutral loss matching and cosine similarity (middle), and neutral loss matching and modified cosine similarity (left). The heat map shows MS/MS spectrum pairs with the corresponding spectrum similarity values as indicated on the axes. Each colored point contains multiple observations out of all 955,228 peptide MS/MS spectrum pairs, as indicated by the color bar. The density plots (gray) show the distributions of the individual similarity measures. (**b**) Spectrum usage (explained intensity) for each of the spectrum similarity measures: Cosine similarity (left), neutral loss matching (middle), and modified cosine similarity (right). Spectrum usage is calculated as the average sum of the relative intensities of all fragments that match in each spectrum of a spectrum pair by the corresponding spectrum similarity measures. (**c**) Evaluation of the spectrum similarities in relation to the relative positions of the modifications. Modification positions are ordered N-terminal (0) to C-terminal (1).

Although pairs of peptide MS/MS spectra can serve as ground truth, because in this case all pairwise modifications are known, peptides are only a subset of molecules. Small molecules are chemically more diverse, and many of the MS/MS spectra will be significantly different. Therefore, we also evaluated the different spectrum similarity measures on a random subset of 10 millions spectrum pairs derived from the GNPS community spectral libraries (**Figure 3**). Because these pairs were randomly selected, the majority of pairs exhibit low spectrum similarities as they correspond to molecules that are structurally unrelated. Similar to the previous analysis using peptide data, the modified cosine similarity strictly outperformed the cosine similarity and neutral loss matching (**Figure 3a**). Neutral loss matching performed considerably better than for peptide data, however, with 32.0% of the spectrum pairs achieving a higher neutral loss match score than cosine score, and 68.0% of the spectrum pairs achieving a higher cosine score than neutral loss score. Because this is a random set of mass spectrum comparisons where most are not expected to match, the spectrum usage is low for all three similarity measures (**Figure 3b**). Nevertheless, the rank of explained intensities, with the explained intensity from the modified cosine similarity outperforming the explained intensity from the cosine similarity, which in turn outperformed the explained intensity from neutral loss matching, was the same as for the peptide data. We also evaluated the spectrum similarity measures in function of the chemical similarity between the molecules, based on the Tanimoto index^23^ as a proxy for structural similarity (**Figure 3c**). This indicated that although even for molecules with a high Tanimoto similarity the majority of pairs still exhibited a poor spectrum similarity, the modified cosine similarity was able to best reflect structural similarity.

**Figure 3.**
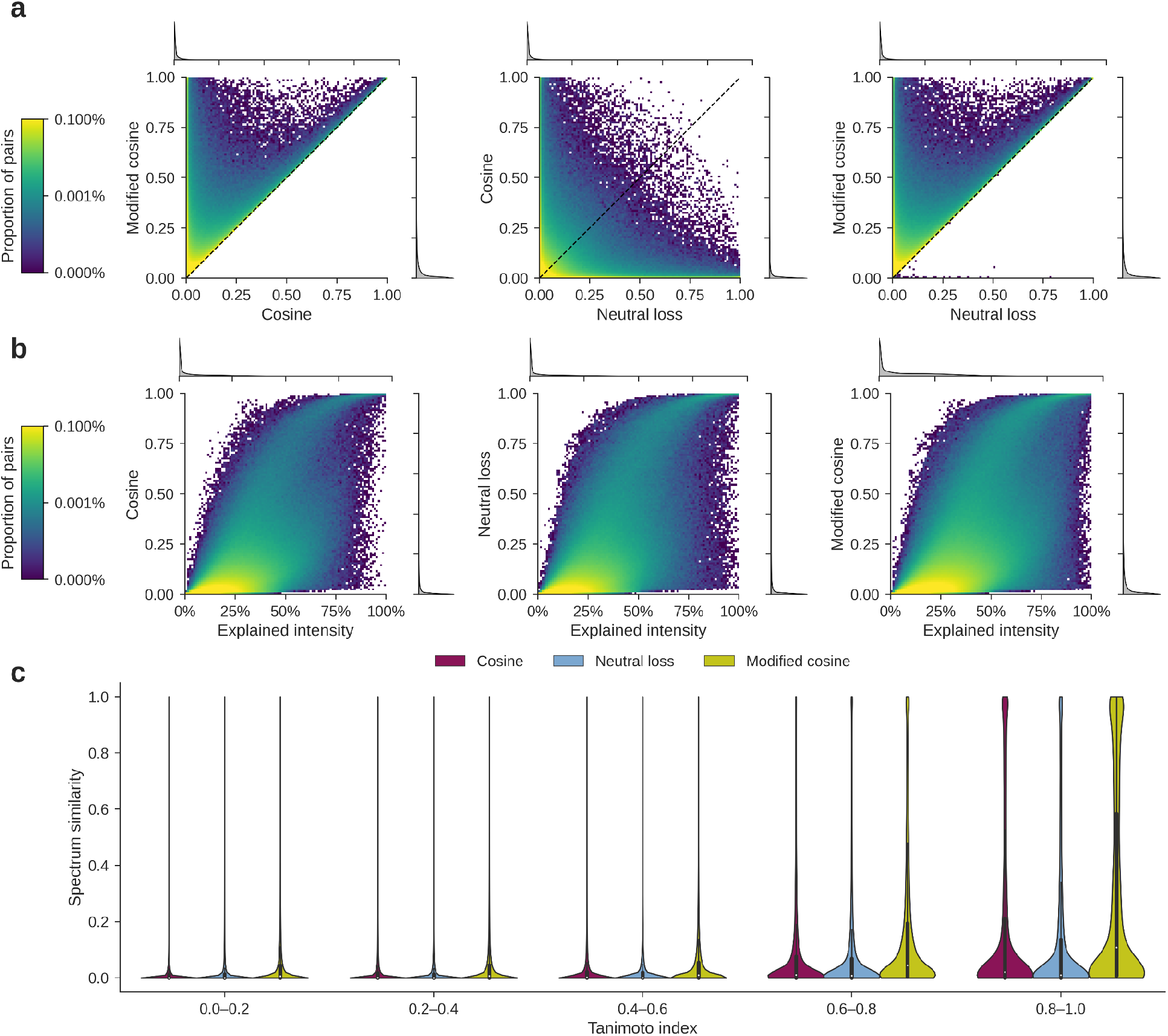
Comparison between cosine similarity, neutral loss matching, and modified cosine similarity when matching small molecule MS/MS spectra. (**a**) Comparison between cosine similarity and modified cosine similarity (left), neutral loss matching and cosine similarity (middle), and neutral loss matching and modified cosine similarity (left). The heat map shows MS/MS spectrum pairs with the corresponding spectrum similarity values as indicated on the axes. Each colored point contains multiple observations out of all 10 million small molecule MS/MS spectrum pairs, as indicated by the color bar. The density plots (gray) show the distributions of the individual similarity measures. (**b**) Spectrum usage (explained intensity) for each of the spectrum similarity measures: Cosine similarity (left), neutral loss matching (right), and modified cosine similarity (right). Spectrum usage is calculated as the average sum of the relative intensities of all fragments that match in each spectrum of a spectrum pair by the corresponding spectrum similarity measures. (**c**) Evaluation of the spectrum similarities in relation to the structural similarities, as measured using the Tanimoto index. A high Tanimoto index (0.8–1) indicates that the pair of molecules has high structural similarity.

Considering that the majority of the 10 million randomly selected spectrum pairs correspond to MS/MS spectra of unrelated molecules, we filtered for mass spectrum pairs whose molecules exhibit high structural similarity (Tanimoto index above 0.9; 8,956 spectrum pairs). Although the Tanimoto index is an imperfect method to fully assess structural similarity,^24^ this filter still enriches for structurally related molecules. This data subset can provide additional insights into the impact of each spectrum similarity measure, especially when comparing the spectrum usage (**Figure 4a**). This indicates that the modified cosine similarity captured more spectrum usage (as defined by explained intensity) than the cosine similarity and neutral loss matching. The different behavior of peptide spectrum pairs versus metabolite spectrum pairs, especially when comparing cosine similarity to neutral loss matching (**Figure 2a, Figure 3a, Figure 4a**) suggests that various modifications may impact spectrum similarity differently. This is supported by the observation that spectrum similarities for structurally related molecules differ significantly per chemical compound class (**Figure 4b**). For example, some compound classes that are characterized by shared or similar molecule backbones, such as purine nucleotides and glycerophospholipids, exhibit very high cosine and modified cosine similarities and low neutral loss scores. In contrast, other compound classes, such as flavonoids, show very low similarities, irrespective of the spectrum similarity measure. Finally, there are interesting compound classes, such as carboxylic acids and indoles, that show lower cosine similarities and neutral loss scores, but higher modified cosine similarities. This indicates that for these classes, the MS/MS spectra contain both directly matching fragment ions and fragment ions that are shifted by modifications or neutral losses, which can only be captured jointly by the modified cosine similarity.

**Figure 4.**
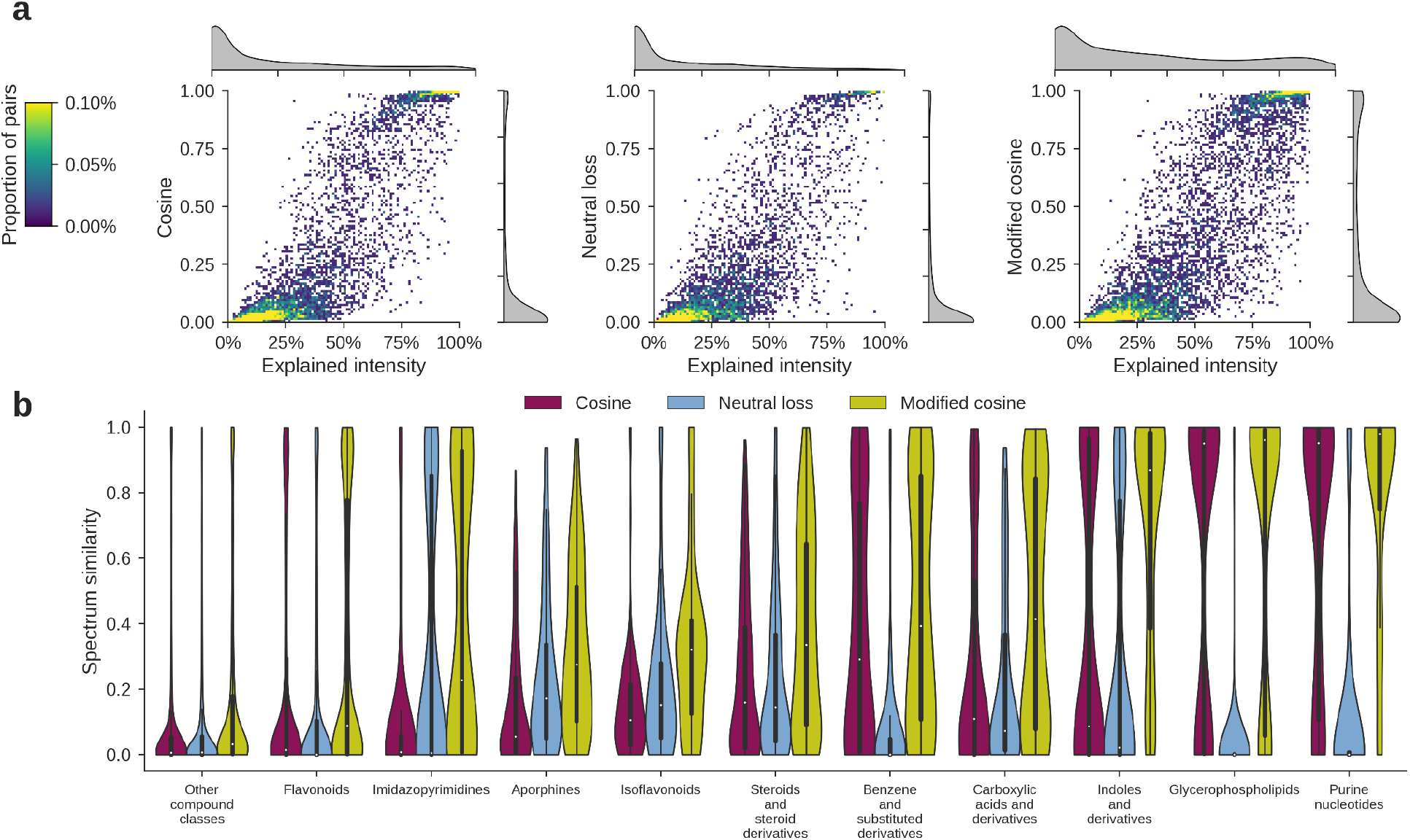
Comparison between cosine similarity, neutral loss matching, and modified cosine similarity when matching structurally similar (Tanimoto index above 0.9) small molecule MS/MS spectra. (**a**) Spectrum usage (explained intensity) for each of the spectrum similarity measures: Cosine similarity (left), neutral loss matching (right), and modified cosine similarity (right). (**b**) Evaluation of the spectrum similarities per compound class, as determined by Classyfire. Violin plots of the similarities for the ten most frequent compound classes are shown, with the remaining compound classes grouped under “Other compound classes.”

To evaluate how specific types of modifications impact spectrum similarity, we performed detailed analyses of the bile acids molecular family, which has a number of distinct modifications. Our lab has recently discovered many new bile acids, for which the MS/MS spectra have been made available as an open access resource.^25–29^ Additionally, we have collected MS/MS data on a historical library of previously determined bile acids. Although other studies have reported the recent discovery of new bile acids as well,^30–35^ unfortunately these could not be included in the current analysis as the corresponding MS/MS data are not publicly available yet. Because our continued study of bile acids has given us a deep understanding of this subset of molecules, we can further analyze these data to understand how specific modifications influence spectrum similarity (**Figure 5**). In total 846 MS/MS spectra of 369 unique bile acids were included, leading to 340,637 pairs of MS/MS spectra. Although bile acids can undergo many different modifications,^36^ here we focused on (i) all bile acid pairs, (ii) bile acid pairs that differ by a single oxygen, (iii) bile acid pairs that differ by a conjugated amino acid, and (iv) bile acids pairs that differ by the substitution of a conjugated amino acid. Similarly as for the previous analyses, the modified cosine similarity strictly outperformed neutral loss matching and the cosine similarity (**Figure 5a**). Interestingly, for this class of molecules neutral loss matching outperformed cosine similarity in 68.6% of cases.

**Figure 5.**
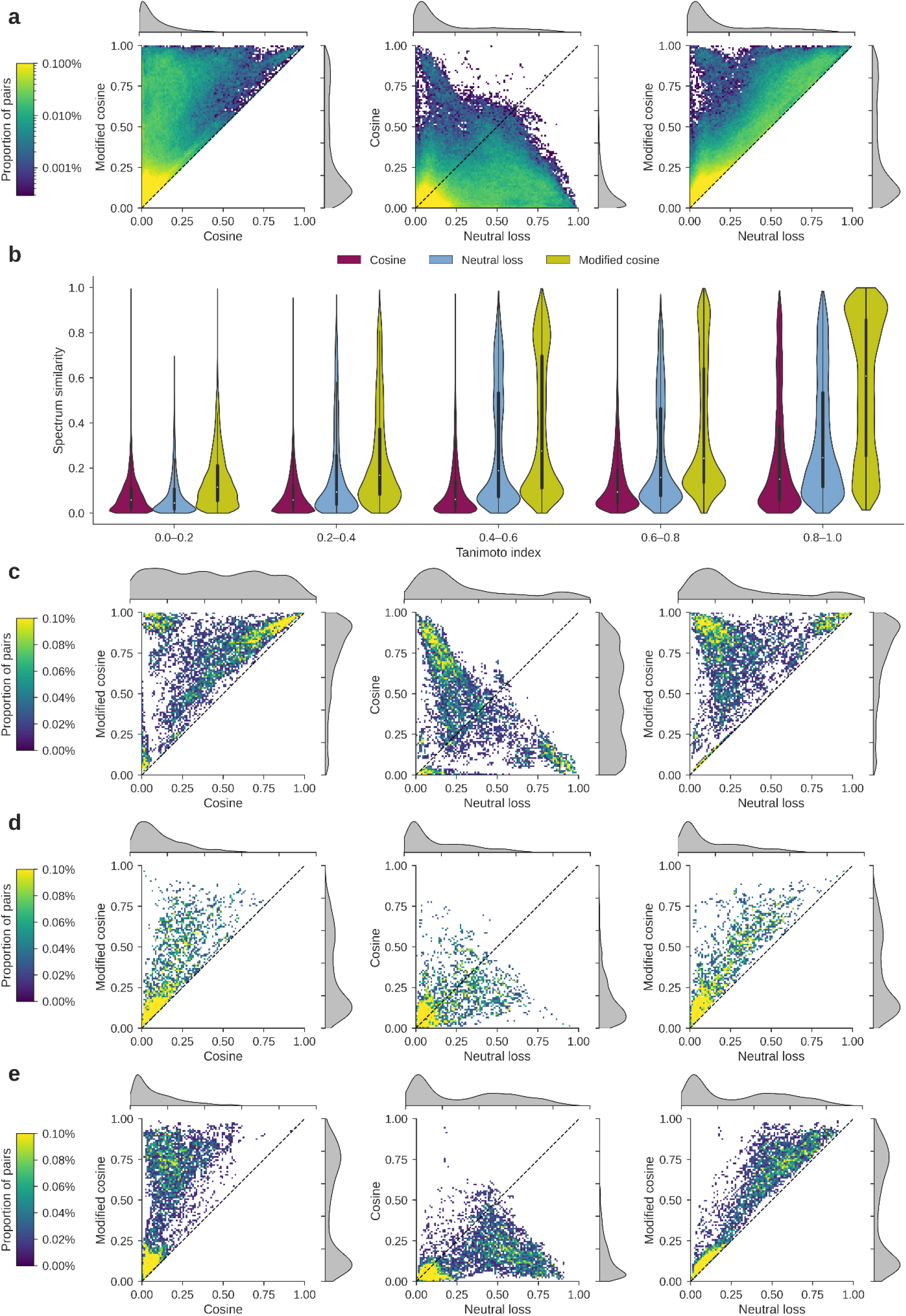
Comparison between cosine similarity, neutral loss matching, and modified cosine similarity when matching MS/MS spectra of bile acids. (**a**) Comparison between cosine similarity and modified cosine similarity (left), neutral loss matching and cosine similarity (middle), and neutral loss matching and modified cosine similarity (left). The heat map shows MS/MS spectrum pairs with the corresponding spectrum similarity values as indicated on the axes. Each colored point contains multiple observations out of all 340,637 bile acid MS/MS spectrum pairs, as indicated by the color bar. The density plots (gray) show the distributions of the individual similarity measures. (**b**) Evaluation of the spectrum similarities in relation to the structural similarities, as measured using the Tanimoto index. A high Tanimoto index (0.8–1) indicates that the pair of molecules has high structural similarity. (**c**) Comparison between spectrum similarity measures for 8,128 bile acid pairs that differ by a single oxygen. (**d**) Comparison between spectrum similarity measures for 3,515 bile acid pairs that differ by a conjugated amino acid. (**e**) Comparison between spectrum similarity measures for 8,624 bile acid pairs that differ by the substitution of a conjugated amino acid.

An increasing Tanimoto index between bile acid pairs, reflecting more similarity in the structures, resulted in increased spectrum similarities as well, with the modified cosine similarity best capturing high structural similarity (**Figure 5b**). However, these trends strongly depend on the type of modification. For a modification consisting of a single oxygen difference, the majority of spectrum pairs achieved high modified cosine similarities (median 0.805), whereas cosine similarities (median 0.426), and neutral loss scores (median 0.224) were considerably lower (**Figure 5c**). For comparing spectra of non-conjugated bile acids to conjugated bile acids, none of the spectrum similarity measures work well (**Figure 5d**). This indicates that this type of modification exerts a strong influence on the fragmentation pathways, despite belonging to the same class of molecules. Finally, when comparing spectra of conjugated bile acids that have undergone an amino acid substitution, neutral loss matching behaved most similarly to the modified cosine similarity, both showing a bimodal score distribution, and strongly outperforming the cosine similarity (**Figure 5e**). Nevertheless, for all of the mass spectrum pairs, there was not a single instance in which neutral loss matching outperformed the modified cosine similarity.

## Discussion

Here we have evaluated the performance of three related mass spectrum similarity measures—cosine similarity, neutral loss matching, and modified cosine similarity—in capturing similarities between structurally related molecules. Our evaluations indicate that the modified cosine similarity is superior to both alternative similarity measures for both peptide and small molecule MS/MS data. These results are in concordance with the popularity of the modified cosine similarity as one of the most commonly used methods to find MS/MS spectra of related molecules.^7^

Our interpretation from the peptide data is that the cosine similarity most closely approximates the modified cosine similarity when the modification is located in the N-terminal region of the peptide, as this will maximally conserve the overall spectral pattern of the dominant y-ions. Furthermore, despite only considering directly matching fragment ions, our results show that the traditional cosine similarity outperforms the recently proposed neutral loss matching strategy^20^ in the majority of cases, even for modified small molecules. Despite the overall advantage of the cosine similarity, there are a significant number of small molecule MS/MS spectrum pairs for which neutral loss matching outperformed cosine similarity. In contrast, neutral loss matching was always inferior to the modified cosine similarity. We currently hypothesize that neutral loss matching outperforms the cosine similarity when the modification is on the same side of the molecule as the charge. In this case, all fragment ions are shifted by the modification mass, and the neutral loss spectra become mirror images of the original MS/MS spectra, whereas no fragment ions will match directly using the cosine similarity. Nevertheless, under such a circumstance the modified cosine similarity still captures all shifted fragment ions as well, which is confirmed by the comparisons presented. In conclusion, both cosine similarity and neutral loss matching can only capture a subset of fragment ions that are matched by the modified cosine similarity.

Although there is not a single test case where our analysis revealed neutral loss matching to outperform the modified cosine similarity, to enable community use of neutral loss matching as a comparative alternative to the cosine similarity and the modified cosine similarity, we have implemented it in several software tools, including matchms,^37^ GNPS,^7^ and MZmine.^38^ For detailed exploration of MS/MS spectrum similarity, user-friendly viewers are also available, for example within MZmine (**Figure 6**).^38,39^ Additionally, to promote open and reproducible science, all data and code to execute the presented analyses are available with an open and unrestricted license.

**Figure 6.**
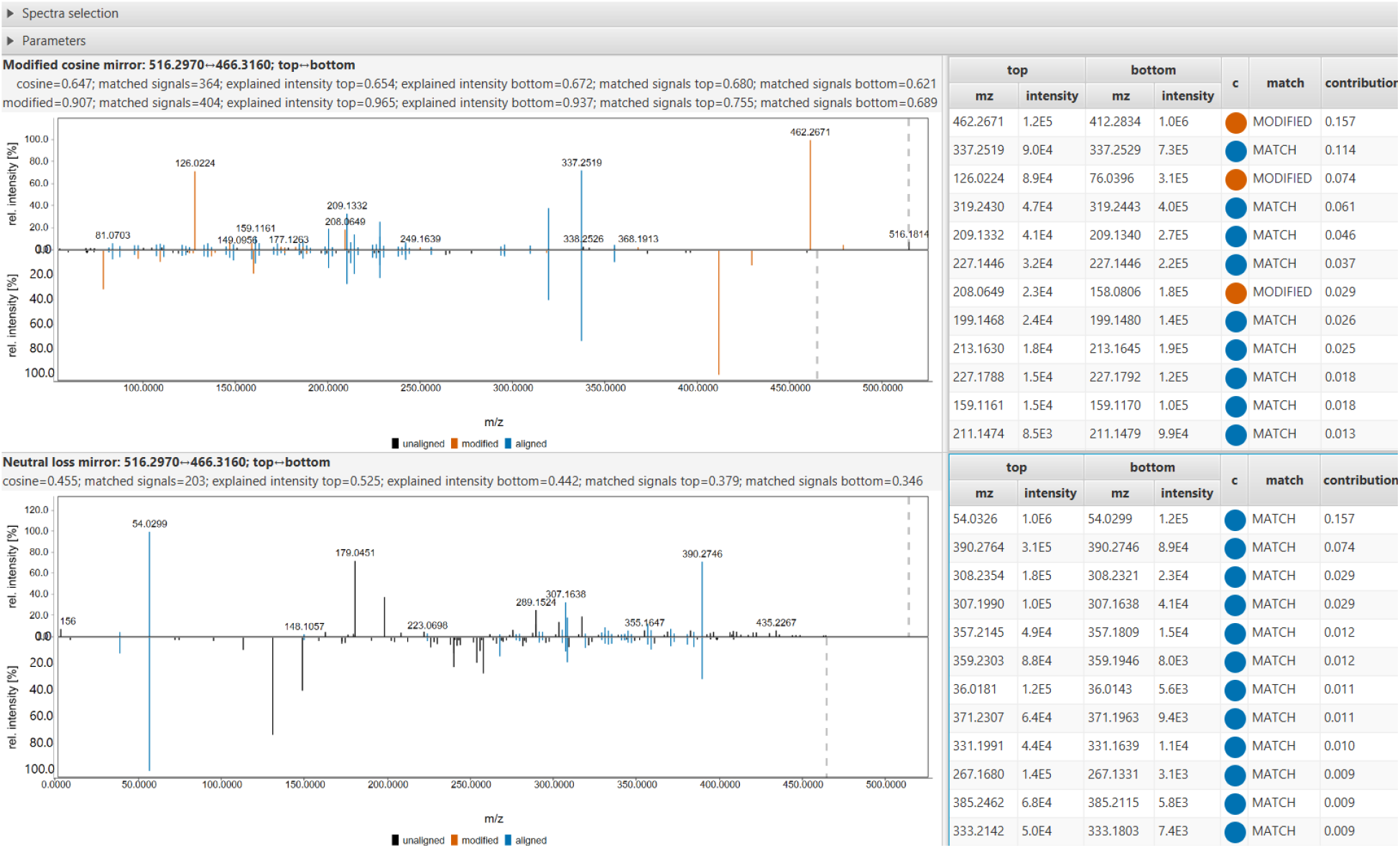
Viewer in MZmine for user-friendly evaluation of spectrum similarity measures. The example shows the spectrum usage from the modified cosine similarity (top) and neutral loss matching (bottom) for the MS/MS spectra of the bile acids taurocholic acid (CCMSLIB00005435561) and glycocholic acid (CCMSLIB00005435513). The viewer shows the directly matching fragment ions (top) and neutral loss ions (bottom) in blue, and the matching fragment ions that are offset by the precursor mass difference (top) in orange. The side panel shows the relative contribution of all pairs of matching fragment and neutral loss ions to the overall score.

Besides the three mass spectrum similarity measures that we have evaluated in this work, an increasing number of other spectrum similarity measures for the discovery of MS/MS spectra of related molecules are being proposed, including various MS/MS spectrum preprocessing steps, different implementations of (modified) cosine similarities that utilize different weighting schemes,^11,19,40^ and methods that use machine learning, such as Spec2Vec,^14^ MS2DeepScore,^16^ spectral entropy,^15^ SIMILE,^17^ GLEAMS,^18^ and many others. These similarity measures likely provide alternative and complementary approaches for the discovery of related molecules, as the modified cosine similarity only accounts for mass shifts associated with a single structural modification, whereas in practice the structural relationship between two analogs can consist of two or more modifications, resulting in more complex mass fragmentation relationships. Future benchmarking of such methods, as well as the effects of data processing, are necessary to provide further insight into the strengths and limitations of each approach.

## Acknowledgments

This research was supported by BBSRC-NSF award 2152526 and National Institutes of Health award U19 AG063744. We want to thank Kris Laukens for assistance with creating the figures. Figure 1 was created with BioRender.com.

## Author contributions statement

PCD conceptualized and supervised the work. WB, RS, and FH implemented various spectrum similarity measures. WB, RS, JJJvdH, MW, and PCD performed the data analyses. MW provided computational resources. WB and PCD wrote the manuscript. All authors reviewed and edited the manuscript.

## Methods

### Spectrum similarity measures

The cosine similarity, modified cosine similarity, and neutral loss matching were implemented in Python (version 3.10). In contrast to the original formulation of neutral loss spectra,^20^ spectra were not mirrored at their precursor *m/z*, as this is not properly defined for multiply charged precursors (i.e. a multiply charged precursor peak can have a lower *m/z* value than its singly charged fragment ions), but instead at the theoretical *m/z* of the singly charged precursor. For the cosine similarity and neutral loss matching, only directly matching fragment ions with a near-identical *m/z* or Δ*m*/*z*, respectively, were considered. For the modified cosine similarity, fragment ions with *m/z* values that are shifted by the pairwise precursor mass difference were considered as well. For all similarity measures, the optimal matching peak assignment across both MS/MS spectra was computed using the SciPy (version 1.8.0)^41^ implementation of the linear assignment problem. Additional scientific computing used Numba (version 0.55.1),^42^ NumPy (version 1.21.5),^43^ and spectrum_utils (version 0.4.0)^44^ for efficient spectrum similarity computations, Matplotlib (version 3.5.1)^45^ and Seaborn (version 0.11.2)^46^ for data visualization, Pyteomics (version 4.5.3)^47^ for data loading, and Jupyter notebooks^48^ and Pandas (version 1.4.1)^49^ for data analysis.

### Analysis of modified peptide MS/MS spectrum pairs

Pairs of modified peptide MS/MS spectra were extracted from the MassIVE-KB spectral library (version 2018/06/15).^21^ This is a repository-wide spectral library derived from over 30 TB of human MS/MS proteomics data, containing 2,154,269 MS/MS spectra. First, MS/MS spectra with precursor charge 2, 3, or 4 were considered and spectrum pairs were required to have identical precursor charges. Second, pairs of spectra whose peptides differ by a single modification were extracted. This modification can consist of the absence/presence of a single post-translational modification, or a single amino acid difference (edit distance of 1, corresponding to a single amino acid substitution or amino acid prefix or suffix addition/removal). To avoid inclusion of MS/MS spectrum pairs that only differ by nearby ambiguously localized post-translational modifications, all MS/MS spectrum pairs were required to have a precursor mass difference of at least 4 *m/z*. This resulted in 955,228 MS/MS spectrum pairs with known peptide labels derived from the MassIVE-KB spectral library.

MS/MS spectra were preprocessed by removing peaks within a 0.1 *m/z* window around the precursor *m/z* and removing noise peaks with intensity below 1% of the base peak intensity. Spectrum similarities were computed using a 0.1 *m/z* fragment mass tolerance. The relative locations of the modifications by which pairs of peptides differ were calculated by normalizing the modification indexes in each pair of peptide sequences by the length of the shortest paired peptide sequence.

### Analysis of modified small molecule MS/MS spectrum pairs

Pairs of modified small molecule MS/MS spectra were extracted from the GNPS community spectral libraries (ALL_GNPS_NO_PROPOGATED), consisting of 495,600 MS/MS spectra, and the GNPS bile acids library (BILELIB19), consisting of 4,533 MS/MS spectra (data downloaded on May 12, 2022).^7^ First, MS/MS spectra with precursor charge 1 were considered (spectra with the unspecified precursor charge 0 were assumed to have precursor charge 1 as well). Additionally, only centroid MS/MS spectra that contain at least 6 fragment ions, were acquired in positive ion mode, and have [M+H]+ adducts were included. For the GNPS community spectral libraries, only MS/MS spectra with structural information specified by InChI or SMILES strings were included, and these were used to compute compound classes using Classyfire via its web application programming interface.^50^

For the general small molecule analysis, all MS/MS spectrum pairs from the GNPS community spectral libraries with a pairwise precursor mass difference between 1 and 200 *m/z* were considered, resulting in over 1.5 billion possible spectrum pairs. In deference to computational limitations, 10 million MS/MS spectrum pairs were randomly selected for the subsequent analysis.

For the modified bile acids analysis, all MS/MS spectrum pairs from the GNPS bile acids library with a pairwise precursor mass difference between 1 and 200 *m/z* were considered, resulting in 340,637 spectrum pairs. Additionally, bile acids that differ by a single oxygen were selected by filtering on a pairwise precursor mass difference (mass tolerance 0.01 *m/z*) of 15.9949 *m/z*; bile acids that differ by a conjugation were selected by filtering on pairwise precursor mass differences (mass tolerance 0.01 *m/z*) of 57.0214 *m/z* (glycine), 71.0371 *m/z* (alanine), 107.0041 *m/z* (taurine), 147.0684 *m/z* (phenylalanine), or 163.0633 *m/z* (tyrosine); and bile acids that differ by a substitution in conjugation were selected by filtering on pairwise precursor mass differences (mass tolerance 0.01 *m/z*) of 49.9826 *m/z* (taurine ↔ glycine), 35.9669 *m/z* (tauro ↔ alanine), 40.0643 *m/z* (taurine ↔ phenylalanine), 56.0592 *m/z* (taurine ↔ tyrosine), 90.0469 *m/z* (glycine ↔ phenylalanine), 106.0418 *m/z* (glycine ↔ tyrosine), 76.0313 *m/z* (alanine ↔ phenylalanine), or 92.0262 *m/z* (alanine ↔ tyrosine).

MS/MS spectra were preprocessed by removing peaks within a 0.1 *m/z* window around the precursor *m/z* and removing noise peaks with intensity below 1% of the base peak intensity. Spectrum similarities were computed using a 0.1 *m/z* fragment mass tolerance. Tanimoto indexes^23^ were computed using RDKit (version 2022.3.2)^51^ by creating molecule objects from their SMILES strings, deriving RDKit topological fingerprints using the RDKit default settings (2048 bits), and computing the Tanimoto index.

### Data and code availability

All code and notebooks to recreate the presented analyses are available as open source under the permissive BSD license at https://github.com/bittremieux/cosine_neutral_loss. A permanent archive of the source code and the analysis notebooks is available on Zenodo at https://doi.org/10.5281/zenodo.6584619. Additionally, cosine similarity, modified cosine similarity, and neutral loss matching have been implemented in the matchms Python package (https://github.com/matchms/matchms; version 0.15.0).^37^ An interactive viewer to inspect neutral loss matching and (modified) cosine similarity is available in MZmine 3 (https://github.com/mzmine/mzmine3)^38^ under “Menu > Tools > Spectral mirror.” Spectra can be retrieved by their GNPS library identifiers or Universal Spectrum Identifiers (USIs).^39,52^ Experimental MS/MS spectra can be selected from feature finding results by selecting two rows in a feature table and choosing “Show > MS/MS spectral mirror” from the right-click context menu.

The MassIVE-KB spectral library and the GNPS community spectral libraries are available under an open and free license at https://massive.ucsd.edu/ProteoSAFe/static/massive-kb-libraries.jsp and https://gnps-external.ucsd.edu/gnpslibrary, respectively. A permanent archive of the spectral libraries, as well as the computed similarity scores for the different analyses, is available on Zenodo at https://doi.org/10.5281/zenodo.6829249.

Individual spectra are accessible by their Universal Spectrum Identifiers (USIs).^39,52^ The spectra displayed in **Figure 1b** and **Figure 6** are:

- Apratoxin A:[mzspec:GNPS:GNPS-LIBRARY:accession:CCMSLIB00000424840](https://metabolomics-usi.ucsd.edu/dashinterface/?usi=mzspec%3AGNPS%3AGNPS-LIBRARY%3Aaccession%3ACCMSLIB00000424840)
- Apratoxin A analog with loss of 30.010 Da: [mzspec:MSV000086109:BD5_dil2x_BD5_01_57213:scan:760](https://metabolomics-usi.ucsd.edu/dashinterface/?usi1=mzspec%3AMSV000086109%3ABD5_dil2x_BD5_01_57213%3Ascan%3A760)
- Taurocholic acid:[mzspec:GNPS:GNPS-LIBRARY:accession:CCMSLIB00005435561](https://metabolomics-usi.ucsd.edu/dashinterface/?usi=mzspec%3AGNPS%3AGNPS-LIBRARY%3Aaccession%3ACCMSLIB00005435561)
- Glycocholic acid:mzspec:[GNPS:GNPS-LIBRARY:accession:CCMSLIB00005435513](https://metabolomics-usi.ucsd.edu/dashinterface/?usi=mzspec%3AGNPS%3AGNPS-LIBRARY%3Aaccession%3ACCMSLIB00005435513)

## For Table of Contents Use Only

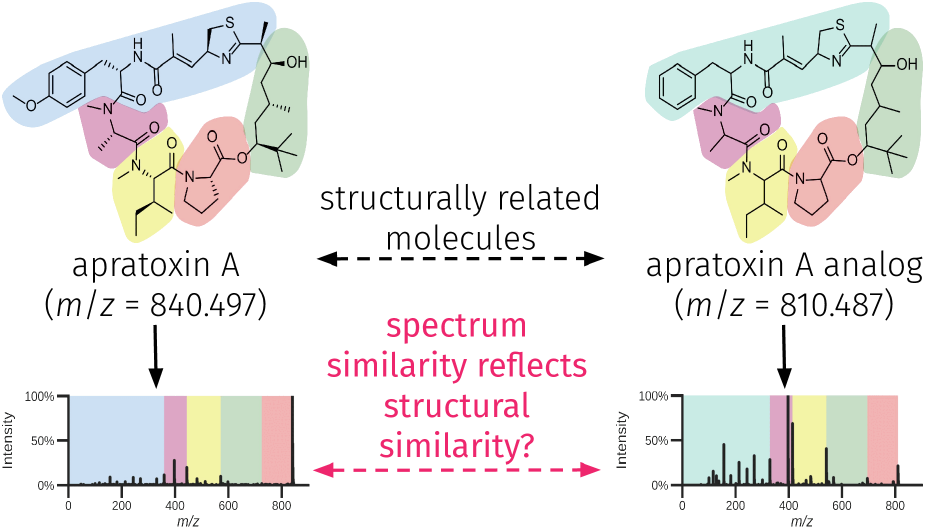

## References

(1) Watrous, J.; Roach, P.; Alexandrov, T.; Heath, B. S.; Yang, J. Y.; Kersten, R. D.; van der Voort, M.; Pogliano, K.; Gross, H.; Raaijmakers, J. M.; Moore, B. S.; Laskin, J.; Bandeira, N.; Dorrestein, P. C. Mass Spectral Molecular Networking of Living Microbial Colonies. Proc. Natl. Acad. Sci. 2012, 109 (26), E1743–E1752. https://doi.org/10.1073/pnas.1203689109.

(2) Stein, S. Mass Spectral Reference Libraries: An Ever-Expanding Resource for Chemical Identification. Anal. Chem. 2012, 84 (17), 7274–7282. https://doi.org/10.1021/ac301205z.

(3) Vinaixa, M.; Schymanski, E. L.; Neumann, S.; Navarro, M.; Salek, R. M.; Yanes, O. Mass Spectral Databases for LC/MS-and GC/MS-Based Metabolomics: State of the Field and Future Prospects. TrAC Trends Anal. Chem. 2016, 78, 23–35. https://doi.org/10.1016/j.trac.2015.09.005.

(4) Beniddir, M. A.; Kang, K. B.; Genta-Jouve, G.; Huber, F.; Rogers, S.; van der Hooft, J. J. J. Advances in Decomposing Complex Metabolite Mixtures Using Substructure-and Network-Based Computational Metabolomics Approaches. Nat. Prod. Rep. 2021, 38 (11), 1967–1993. https://doi.org/10.1039/D1NP00023C.

(5) Stein, S. E.; Scott, D. R. Optimization and Testing of Mass Spectral Library Search Algorithms for Compound Identification. J. Am. Soc. Mass Spectrom. 1994, 5 (9), 859–866. https://doi.org/10.1016/1044-0305(94)87009-8.

(6) Wan, K. X.; Vidavsky, I.; Gross, M. L. Comparing Similar Spectra: From Similarity Index to Spectral Contrast Angle. J. Am. Soc. Mass Spectrom. 2002, 13 (1), 85–88. https://doi.org/10.1016/S1044-0305(01)00327-0.

(7) Wang, M.; Carver, J. J.; Phelan, V. V.; Sanchez, L. M.; Garg, N.; Peng, Y.; Nguyen, D. D.; Watrous, J.; Kapono, C. A.; Luzzatto-Knaan, T.; Porto, C.; Bouslimani, A.; Melnik, A. V.; Meehan, M. J.; Liu, W.-T.; Crüsemann, M.; Boudreau, P. D.; Esquenazi, E.; Sandoval-Calderón, M.; Kersten, R. D.; Pace, L. A.; Quinn, R. A.; Duncan, K. R.; Hsu, C.-C.; Floros, D. J.; Gavilan, R. G.; Kleigrewe, K.; Northen, T.; Dutton, R. J.; Parrot, D.; Carlson, E. E.; Aigle, B.; Michelsen, C. F.; Jelsbak, L.; Sohlenkamp, C.; Pevzner, P.; Edlund, A.; McLean, J.; Piel, J.; Murphy, B. T.; Gerwick, L.; Liaw, C.-C.; Yang, Y.-L.; Humpf, H.-U.; Maansson, M.; Keyzers, R. A.; Sims, A. C.; Johnson, A. R.; Sidebottom, A. M.; Sedio, B. E.; Klitgaard, A.; Larson, C. B.; Boya P, C. A.; Torres-Mendoza, D.; Gonzalez, D. J.; Silva, D. B.; Marques, L. M.; Demarque, D. P.; Pociute, E.; O’Neill, E. C.; Briand, E.; Helfrich, E. J. N.; Granatosky, E. A.; Glukhov, E.; Ryffel, F.; Houson, H.; Mohimani, H.; Kharbush, J. J.; Zeng, Y.; Vorholt, J. A.; Kurita, K. L.; Charusanti, P.; McPhail, K. L.; Nielsen, K. F.; Vuong, L.; Elfeki, M.; Traxler, M. F.; Engene, N.; Koyama, N.; Vining, O. B.; Baric, R.; Silva, R. R.; Mascuch, S. J.; Tomasi, S.; Jenkins, S.; Macherla, V.; Hoffman, T.; Agarwal, V.; Williams, P. G.; Dai, J.; Neupane, R.; Gurr, J.; Rodríguez, A. M. C.; Lamsa, A.; Zhang, C.; Dorrestein, K.; Duggan, B. M.; Almaliti, J.; Allard, P.-M.; Phapale, P.; Nothias, L.-F.; Alexandrov, T.; Litaudon, M.; Wolfender, J.-L.; Kyle, J. E.; Metz, T. O.; Peryea, T.; Nguyen, D.-T.; VanLeer, D.; Shinn, P.; Jadhav, A.; Müller, R.; Waters, K. M.; Shi, W.; Liu, X.; Zhang, L.; Knight, R.; Jensen, P. R.; Palsson, B. Ø.; Pogliano, K.; Linington, R. G.; Gutiérrez, M.; Lopes, N. P.; Gerwick, W. H.; Moore, B. S.; Dorrestein, P. C.; Bandeira, N. Sharing and Community Curation of Mass Spectrometry Data with Global Natural Products Social Molecular Networking. Nat. Biotechnol. 2016, 34 (8), 828–837. https://doi.org/10.1038/nbt.3597.

(8) Moorthy, A. S.; Wallace, W. E.; Kearsley, A. J.; Tchekhovskoi, D. V.; Stein, S. E. Combining Fragment-Ion and Neutral-Loss Matching during Mass Spectral Library Searching: A New General Purpose Algorithm Applicable to Illicit Drug Identification. Anal. Chem. 2017, 89 (24), 13261–13268. https://doi.org/10.1021/acs.analchem.7b03320.

(9) Aron, A. T.; Gentry, E. C.; McPhail, K. L.; Nothias, L.-F.; Nothias-Esposito, M.; Bouslimani, A.; Petras, D.; Gauglitz, J. M.; Sikora, N.; Vargas, F.; van der Hooft, J. J. J.; Ernst, M.; Kang, K. B.; Aceves, C. M.; Caraballo-Rodríguez, A. M.; Koester, I.; Weldon, K. C.; Bertrand, S.; Roullier, C.; Sun, K.; Tehan, R. M.; Boya P, C. A.; Christian, M. H.; Gutiérrez, M.; Ulloa, A. M.; Tejeda Mora, J. A.; Mojica-Flores, R.; Lakey-Beitia, J.; Vásquez-Chaves, V.; Zhang, Y.; Calderón, A. I.; Tayler, N.; Keyzers, R. A.; Tugizimana, F.; Ndlovu, N.; Aksenov, A. A.; Jarmusch, A. K.; Schmid, R.; Truman, A. W.; Bandeira, N.; Wang, M.; Dorrestein, P. C. Reproducible Molecular Networking of Untargeted Mass Spectrometry Data Using GNPS. Nat. Protoc. 2020, 15 (6), 1954–1991. https://doi.org/10.1038/s41596-020-0317-5.

(10) Bandeira, N.; Tsur, D.; Frank, A.; Pevzner, P. A. Protein Identification by Spectral Networks Analysis. Proc. Natl. Acad. Sci. 2007, 104 (15), 6140–6145. https://doi.org/10.1073/pnas.0701130104.

(11) Burke, M. C.; Mirokhin, Y. A.; Tchekhovskoi, D. V.; Markey, S. P.; Heidbrink Thompson, J.; Larkin, C.; Stein, S. E. The Hybrid Search: A Mass Spectral Library Search Method for Discovery of Modifications in Proteomics. J. Proteome Res. 2017. https://doi.org/10.1021/acs.jproteome.6b00988.

(12) Bittremieux, W.; Meysman, P.; Noble, W. S.; Laukens, K. Fast Open Modification Spectral *Library Searching through Approximate Nearest Neighbor Indexing*. J. Proteome Res. 2018, 17 (10), 3463–3474. https://doi.org/10.1021/acs.jproteome.8b00359.

(13) Bittremieux, W.; Laukens, K.; Noble, W. S. Extremely Fast and Accurate Open Modification Spectral Library Searching of High-Resolution Mass Spectra Using Feature Hashing and Graphics Processing Units. J. Proteome Res. 2019, 18 (10), 3792–3799. https://doi.org/10.1021/acs.jproteome.9b00291.

(14) Huber, F.; Ridder, L.; Verhoeven, S.; Spaaks, J. H.; Diblen, F.; Rogers, S.; van der Hooft, J. J. J. Spec2Vec: Improved Mass Spectral Similarity Scoring through Learning of Structural Relationships. PLOS Comput. Biol. 2021, 17 (2), e1008724. https://doi.org/10.1371/journal.pcbi.1008724.

(15) Li, Y.; Kind, T.; Folz, J.; Vaniya, A.; Mehta, S. S.; Fiehn, O. Spectral Entropy Outperforms *MS/MS Dot Product Similarity for Small-Molecule Compound Identification*. Nat. Methods 2021, 18 (12), 1524–1531. https://doi.org/10.1038/s41592-021-01331-z.

(16) Huber, F.; van der Burg, S.; van der Hooft, J. J. J.; Ridder, L. MS2DeepScore: A Novel Deep Learning Similarity Measure to Compare Tandem Mass Spectra. J. Cheminformatics 2021, 13 (1), 84. https://doi.org/10.1186/s13321-021-00558-4.

(17) Treen, D. G. C.; Wang, M.; Xing, S.; Louie, K. B.; Huan, T.; Dorrestein, P. C.; Northen, T. R.; Bowen, B. P. SIMILE Enables Alignment of Tandem Mass Spectra with Statistical Significance. Nat. Commun. 2022, 13 (1), 2510. https://doi.org/10.1038/s41467-022-30118-9.

(18) Bittremieux, W.; May, D. H.; Bilmes, J.; Noble, W. S. A Learned Embedding for Efficient Joint Analysis of Millions of Mass Spectra. Nat. Methods 2022, in press. https://doi.org/10.1101/483263.

(19) Xue, J.; Guijas, C.; Benton, H. P.; Warth, B.; Siuzdak, G. METLIN MS2 Molecular Standards Database: A Broad Chemical and Biological Resource. Nat. Methods 2020, 17 (10), 953–954. https://doi.org/10.1038/s41592-020-0942-5.

(20) Aisporna, A.; Benton, H. P.; Chen, A.; Derks, R. J. E.; Galano, J. M.; Giera, M.; Siuzdak, G. Neutral Loss Mass Spectral Data Enhances Molecular Similarity Analysis in METLIN. J. Am. Soc. Mass Spectrom. 2022, 33 (3), 530–534. https://doi.org/10.1021/jasms.1c00343.

(21) Wang, M.; Wang, J.; Carver, J.; Pullman, B. S.; Cha, S. W.; Bandeira, N. Assembling the Community-Scale Discoverable Human Proteome. Cell Syst. 2018, 7 (4), 412–421.e5. https://doi.org/10.1016/j.cels.2018.08.004.

(22) Bittremieux, W.; Avalon, N. E.; Thomas, S. P.; Kakhkhorov, S. A.; Aksenov, A. A.; Gomes, P. W. P.; Aceves, C. M.; Caraballo Rodriguez, A. M.; Gauglitz, J. M.; Gerwick, W. H.; Jarmusch, A. K.; Kaddurah-Daouk, R. F.; Kang, K. B.; Kim, H. W.; Kondic, T; Mannochio-Russo, H.; Meehan, M. J.; Melnik, A.; Nothias, L.-F.; O’Donovan, C.; Panitchpakdi, M.; Petras, D.; Schmid, R.; Schymanski, E. L.; van der Hooft, J. J. J.; Weldon, K. C.; Yang, H.; Zemlin, J.; Wang, M.; Dorrestein, P. C. Open Access Repository-Scale Propagated Nearest Neighbor Suspect Spectral Library for Untargeted Metabolomics. bioRxiv 2022. https://doi.org/10.1101/2022.05.15.490691.

(23) Tanimoto, T. T. An Elementary Mathematical Theory of Classification and Prediction; International Business Machines Corp., 1958.

(24) Bajusz, D.; Rácz, A.; Héberger, K. Why Is Tanimoto Index an Appropriate Choice for Fingerprint-Based Similarity Calculations? J. Cheminformatics 2015, 7 (1), 20. https://doi.org/10.1186/s13321-015-0069-3.

(25) Quinn, R. A.; Melnik, A. V.; Vrbanac, A.; Fu, T.; Patras, K. A.; Christy, M. P.; Bodai, Z.; Belda-Ferre, P.; Tripathi, A.; Chung, L. K.; Downes, M.; Welch, R. D.; Quinn, M.; Humphrey, G.; Panitchpakdi, M.; Weldon, K. C.; Aksenov, A.; da Silva, R.; Avila-Pacheco, J.; Clish, C.; Bae, S.; Mallick, H.; Franzosa, E. A.; Lloyd-Price, J.; Bussell, R.; Thron, T.; Nelson, A. T.; Wang, M.; Leszczynski, E.; Vargas, F.; Gauglitz, J. M.; Meehan, M. J.; Gentry, E.; Arthur, T. D.; Komor, A. C.; Poulsen, O.; Boland, B. S.; Chang, J. T.; Sandborn, W. J.; Lim, M.; Garg, N.; Lumeng, J. C.; Xavier, R. J.; Kazmierczak, B. I.; Jain, R.; Egan, M.; Rhee, K. E.; Ferguson, D.; Raffatellu, M.; Vlamakis, H.; Haddad, G. G.; Siegel, D.; Huttenhower, C.; Mazmanian, S. K.; Evans, R. M.; Nizet, V.; Knight, R.; Dorrestein, P. C. Global Chemical Effects of the Microbiome Include New Bile-Acid Conjugations. Nature 2020, 579 (7797), 123–129. https://doi.org/10.1038/s41586-020-2047-9.

(26) Dethloff, F.; Vargas, F.; Elijah, E.; Quinn, R.; Park, D. I.; Herzog, D. P.; Müller, M. B.; Gentry, E. C.; Knight, R.; Gonzalez, A.; Dorrestein, P. C.; Turck, C. W. Paroxetine Administration Affects Microbiota and Bile Acid Levels in Mice. Front. Psychiatry 2020, 11, 518. https://doi.org/10.3389/fpsyt.2020.00518.

(27) Petras, D.; Caraballo-Rodríguez, A. M.; Jarmusch, A. K.; Molina-Santiago, C.; Gauglitz, J. M.; Gentry, E. C.; Belda-Ferre, P.; Romero, D.; Tsunoda, S. M.; Dorrestein, P. C.; Wang, M. Chemical Proportionality within Molecular Networks. Anal. Chem. 2021, 93 (38), 12833–12839. https://doi.org/10.1021/acs.analchem.1c01520.

(28) Hoffmann, M. A.; Nothias, L.-F.; Ludwig, M.; Fleischauer, M.; Gentry, E. C.; Witting, M.; Dorrestein, P C.; Dührkop, K.; Böcker, S. High-Confidence Structural Annotation of Metabolites Absent from Spectral Libraries. Nat. Biotechnol. 2021, 40 (3), 411–421. https://doi.org/10.1038/s41587-021-01045-9.

(29) Gentry, E.; Collins, S.; Panitchpakdi, M.; Belda-Ferre, P.; Stewart, A.; Wang, M.; Jarmusch, A.; Avila-Pacheco, J.; Plichta, D.; Aron, A.; Vlamakis, H.; Ananthakrishnan, A.; Clish, C.; Xavier, R.; Baker, E.; Patterson, A.; Knight, R.; Siegel, D.; Dorrestein, P. C. A Synthesis-Based Reverse Metabolomics Approach for the Discovery of Chemical Structures from Humans and Animals. Res. Sq. 2021. https://doi.org/10.21203/rs.3.rs-820302/v1.

(30) Lucas, L. N.; Barrett, K.; Kerby, R. L.; Zhang, Q.; Cattaneo, L. E.; Stevenson, D.; Rey, F. E.; Amador-Noguez, D. Dominant Bacterial Phyla from the Human Gut Show Widespread Ability to Transform and Conjugate Bile Acids. mSystems 2021, 6 (4), e00805–21. https://doi.org/10.1128/mSystems.00805-21.

(31) Zhu, Q.-F.; Wang, Y.-Z.; An, N.; Hao, J.-D.; Mei, P.-C.; Bai, Y.-L.; Hu, Y.-N.; Bai, P.-R.; Feng, Y.-Q. Alternating Dual-Collision Energy Scanning Mass Spectrometry Approach: Discovery of Novel Microbial Bile-Acid Conjugates. Anal. Chem. 2022, 94 (5), 2655–2664. https://doi.org/10.1021/acs.analchem.1c05272.

(32) Garcia, C. J.; Kosek, V.; Beltrán, D.; Tomás-Barberán, F. A.; Hajslova, J. Production of New Microbially Conjugated Bile Acids by Human Gut Microbiota. Biomolecules 2022, 12 (5), 687. https://doi.org/10.3390/biom12050687.

(33) Ma, Y.; Cao, Y.; Song, X.; Zhang, Y.; Li, J.; Wang, Y.; Wu, X.; Qi, X. BAFinder: A Software for Unknown Bile Acid Identification Using Accurate Mass LC-MS/MS in Positive and Negative Modes. Anal. Chem. 2022, 94 (16), 6242–6250. https://doi.org/10.1021/acs.analchem.1c05648.

(34) Shalon, D.; Culver, R. N.; Grembi, J. A.; Folz, J.; Treit, P.; Dethlefsen, L.; Meng, X.; Yaffe, E.; Spencer, S.; Shi, H.; Aranda-Díaz, A.; Patterson, A. D.; Triadafilopoulos, G.; Holmes, S. P.; Mann, M.; Fiehn, O.; Relman, D. A.; Huang, K. C. Profiling of the Human Intestinal Microbiome and Bile Acids under Physiologic Conditions Using an Ingestible Sampling Device. bioRxiv 2022. https://doi.org/10.1101/2022.01.19.476920.

(35) Neugebauer, K. A.; Guzior, D. V.; Feiner, J.; Rzepka, M.; Schillmiller, A.; O’Reilly, S.; Jones, A. D.; Watson, V. E.; Luyendyk, J. P.; McCabe, L.; Quinn, R. A. Bile Acid-CoA:Amino Acid N-Acyltransferase Gene Knockout Alters Early Life Development, the Gut Microbiome and Reveals Unusual Bile Acid Conjugates in Mice. bioRxiv 2022. https://doi.org/10.1101/2022.04.10.487642.

(36) Hofmann, A. F.; Hagey, L. R. Key Discoveries in Bile Acid Chemistry and Biology and Their Clinical Applications: History of the Last Eight Decades. J. Lipid Res. 2014, 55 (8), 1553–1595. https://doi.org/10.1194/jlr.R049437.

(37) Huber, F.; Verhoeven, S.; Meijer, C.; Spreeuw, H.; Castilla, E.; Geng, C.; van der Hooft, J.; Rogers, S.; Belloum, A.; Diblen, F.; Spaaks, J. Matchms - Processing and Similarity Evaluation of Mass Spectrometry Data. J. Open Source Softw. 2020, 5 (52), 2411. https://doi.org/10.21105/joss.02411.

(38) Pluskal, T.; Castillo, S.; Villar-Briones, A.; Orešič, M. MZmine 2: Modular Framework for Processing, Visualizing, and Analyzing Mass Spectrometry-Based Molecular Profile Data. BMC Bioinf. 2010, 11 (1), 395. https://doi.org/10.1186/1471-2105-11-395.

(39) Bittremieux, W.; Chen, C.; Dorrestein, P. C.; Schymanski, E. L.; Schulze, T.; Neumann, S.; Meier, R.; Rogers, S.; Wang, M. Universal MS/MS Visualization and Retrieval with the Metabolomics Spectrum Resolver Web Service. bioRxiv 2020. https://doi.org/10.1101/2020.05.09.086066.

(40) Chen, L.; Lu, W.; Wang, L.; Xing, X.; Chen, Z.; Teng, X.; Zeng, X.; Muscarella, A. D.; Shen, Y.; Cowan, A.; McReynolds, M. R.; Kennedy, B. J.; Lato, A. M.; Campagna, S. R.; Singh, M.; Rabinowitz, J. D. Metabolite Discovery through Global Annotation of Untargeted Metabolomics Data. Nat. Methods 2021, 18 (11), 1377–1385. https://doi.org/10.1038/s41592-021-01303-3.

(41) SciPy 1.0 Contributors; Virtanen, P.; Gommers, R.; Oliphant, T. E.; Haberland, M.; Reddy, T.; Cournapeau, D.; Burovski, E.; Peterson, P.; Weckesser, W.; Bright, J.; van der Walt, S. J.; Brett, M.; Wilson, J.; Millman, K. J.; Mayorov, N.; Nelson, A. R. J.; Jones, E.; Kern, R.; Larson, E.; Carey, C. J.; Polat, i.; Feng, Y.; Moore, E. W.; VanderPlas, J.; Laxalde, D.; Perktold, J.; Cimrman, R.; Henriksen, I.; Quintero, E. A.; Harris, C. R.; Archibald, A. M.; Ribeiro, A. H.; Pedregosa, F; van Mulbregt, P. SciPy 1.0: Fundamental Algorithms for Scientific Computing in Python. Nat. Methods 2020. https://doi.org/10.1038/s41592-019-0686-2.

(42) Lam, S. K.; Pitrou, A.; Seibert, S. Numba: A LLVM-Based Python JIT Compiler. In Proceedings of the Second Workshop on the LLVM Compiler Infrastructure in HPC - LLVM ‘15; ACM Press: Austin, TX, USA, 2015; pp 1–6. https://doi.org/10.1145/2833157.2833162.

(43) Harris, C. R.; Millman, K. J.; van der Walt, S. J.; Gommers, R.; Virtanen, P.; Cournapeau, D.; Wieser, E.; Taylor, J.; Berg, S.; Smith, N. J.; Kern, R.; Picus, M.; Hoyer, S.; van Kerkwijk, M. H.; Brett, M.; Haldane, A.; del Río, J. F.; Wiebe, M.; Peterson, P.; Gérard-Marchant, P.; Sheppard, K.; Reddy, T.; Weckesser, W.; Abbasi, H.; Gohlke, C.; Oliphant, T. E. Array Programming with NumPy. Nature 2020, 585 (7825), 357–362. https://doi.org/10.1038/s41586-020-2649-2.

(44) Bittremieux, W. spectrum_utils: A Python Package for Mass Spectrometry Data Processing and Visualization. Anal. Chem. 2020, 92 (1), 659–661. https://doi.org/10.1021/acs.analchem.9b04884.

(45) Hunter, J. D. Matplotlib: A 2D Graphics Environment. Comput. Sci. Eng. 2007, 9 (3), 90–95. https://doi.org/10.1109/MCSE.2007.55.

(46) Waskom, M. L. Seaborn: Statistical Data Visualization. J. Open Source Softw. 2021, 6 (60), 3021. https://doi.org/10.21105/joss.03021.

(47) Levitsky, L. I.; Klein, J. A.; Ivanov, M. V.; Gorshkov, M. Pyteomics 4.0: Five Years of Development of a Python Proteomics Framework. J. Proteome Res. 2019, 18 (2), 709–714. https://doi.org/10.1021/acs.jproteome.8b00717.

(48) Thomas, K.; Benjamin, R.-K.; Fernando, P.; Brian, G.; Matthias, B.; Jonathan, F.; Kyle, K.; Jessica, H.; Jason, G.; Sylvain, C.; Paul, I.; Damián, A.; Safia, A.; Carol, W.; Jupyter Development Team. Jupyter Notebooks -- A Publishing Format for Reproducible Computational Workflows. In Positioning and Power in Academic Publishing: Players, Agents and Agendas; IOS Press, 2016; pp 87–90.

(49) McKinney, W. Data Structures for Statistical Computing in Python. In Proceedings of the 9th Python in Science Conference; van der Walt, S., Millman, J., Eds.; Austin, Texas, USA, 2010; pp 51–56.

(50) Djoumbou Feunang, Y.; Eisner, R.; Knox, C.; Chepelev, L.; Hastings, J.; Owen, G.; Fahy, E.; Steinbeck, C.; Subramanian, S.; Bolton, E.; Greiner, R.; Wishart, D. S. ClassyFire: *Automated Chemical Classification with a Comprehensive, Computable Taxonomy*. J. Cheminformatics 2016, 8 (1), 61. https://doi.org/10.1186/s13321-016-0174-y.

(51) Landrum, G.; Tosco, P.; Kelley, B.; Ric; Sriniker; Gedeck; Vianello, R.; NadineSchneider; Kawashima, E.; Dalke, A.; N, D.; Cosgrove, D.; Cole, B.; Swain, M.; Turk, S.; AlexanderSavelyev; Jones, G.; Vaucher, A.; Wójcikowski, M.; Ichiru Take; Probst, D.; Ujihara, K.; Scalfani, V. F.; Godin, G.; Pahl, A.; Francois Berenger; JLVarjo; Strets123; JP; DoliathGavid. Rdkit/Rdkit: 2022_O3_2 (Q1 2022) Release; Zenodo, 2022. https://doi.org/10.5281/ZENODO.6483170.

(52) Deutsch, E. W.; Perez-Riverol, Y.; Carver, J.; Kawano, S.; Mendoza, L.; Van Den Bossche, T.; Gabriels, R.; Binz, P.-A.; Pullman, B.; Sun, Z.; Shofstahl, J.; Bittremieux, W.; Mak, T.; Klein, J.; Zhu, Y.; Lam, H.; Vizcaino, J. A.; Bandeira, N. Universal Spectrum Identifier for Mass Spectra. Nat. Methods 2021, 18, 768–770. https://doi.org/10.1038/s41592-021-01184-6.

